# Identification of a novel caspase cleavage site in huntingtin that regulates mutant huntingtin clearance

**DOI:** 10.1101/232827

**Authors:** D.D.O. Martin, M. E. Schmidt, Y. T. Nguyen, N. Lazic, M. R. Hayden

**Author notes:** Corresponding author: Dr. Michael R. Hayden.

## Abstract

Huntington disease (HD) is a progressive neurodegenerative disease that initially affects the striatum leading to changes in behavior and loss of motor coordination. It is caused by an expansion in the polyglutamine repeat at the N-terminus of huntingtin (HTT) that leads to aggregation of mutant HTT. The loss of wildtype function, in combination with the toxic gain of function mutation, initiates various cell death pathways. Wildtype and mutant HTT are regulated by different post-translational modifications that can positively or negatively regulate their function or toxicity. In particular, we have previously shown that caspase cleavage of mutant HTT at amino acid position aspartate 586 (D586) by caspase-6 is critical for the pathogenesis of the disease in an HD mouse model. Herein, we describe the identification of a new caspase cleavage site at position D572 that is mediated by caspase-1. Inhibition of caspase-1 also inhibits cleavage at D586 through inhibition of caspase-6. Inhibition of caspase cleavage at D572 significantly decreases mutant HTT aggregation and significantly increased the turnover of soluble mutant HTT. This suggests that caspase-1 may be a viable target to inhibit caspase cleavage of mutant HTT at both D572 and D586 to promote mutant HTT clearance.

## Introduction

Huntington disease (HD) is a devastating neurological disease that is characterised by progressive neuronal degeneration, particularly in the striatum, leading to alterations in behavior and loss of motor coordination. There is currently no treatment to delay progression of the disease. HD is a monogenic dominantly inherited disease caused by an expansion of a CAG repeat in the huntingtin (*HTT*) gene that encodes for an extended polyglutamine in the N-terminus of the HTT protein (1). The expanded polyglutamine leads to a toxic gain of function partly due to protein aggregation of mutant HTT (mHTT) that is compounded by the toxic loss of function of wildtype HTT (wtHTT), which has been shown to be involved in many cellular processes including intracellular trafficking (2), autophagy (3, 4), and cell spindle assembly (5).

A key step in the pathogenesis of HD is mediated by proteolysis of HTT, predominantly by caspases. Several sites of cleavage have been identified, including D513, D552 and D586 (6–8). Previously, we have shown that proteolysis of mHTT at D586 is critical for the disease progression in mice (8). Caspase cleavage at D586 is primarily mediated by caspase-6, and to a lesser degree caspase-8 (9). Of note, blocking caspase cleavage at D586 completely ameliorates the HD phenotype in the YAC HD mouse model (8). HTT has also been shown to be cleaved by caspases at D552 and D513 (6–8). Cleavage at D513 is considered toxic, particularly when coupled with calpain cleavage at R167 (6). Interestingly, we have shown that cleavage at D552 is required to release a newly exposed N-terminal glycine position at 553 (G553) that is subsequently post-translationally myristoylated (10–12). Myristoylation involves the covalent addition of the saturated fatty acid myristate to N-terminal glycines through an amide bond that directs proteins to membranes (13, 14). We also recently discovered a naturally occurring missense mutation at this site wherein the essential glycine is substituted to a glutamate, thereby blocking myristoylation (11). Surprisingly, this mutation also blocked caspase cleavage at D552 while also promoting caspase cleavage at D513. Although this mutation was restricted to the control allele, it was found to be toxic when overexpressed in cells.

This suggests an interplay among the HTT post-translational modification (PTM) and caspase networks and that alterations at one site may regulate several downstream PTMs. In addition, we also noted an unknown proteolytic fragment by SDS-PAGE that was increased after the induction of apoptosis, particularly when proteolysis at D586 was blocked. Herein, we describe the identification of this previously unknown band as a novel caspase cleavage site (D572) that regulates both mHTT aggregation and clearance.

## Materials and Methods

### Materials

Click chemistry and cell culture reagents were purchased from Invitrogen (Carlsbad, CA, USA). Affinity purified goat and rabbit anti-GFP antibodies were acquired from Eusera (Edmonton, AB, Canada) and gBlocks for cloning were obtained from IDT (Canada). Caspase inhibitors and purified caspases were purchased from EMD Millipore and Enzo Life Sciences, respectively.

### Cloning

All point mutations were generated using gBlocks, as previously described (9), and HTT_1-588_-YFP (15). Briefly, gBlocks containing the indicated point mutation were generated (IDT) and inserted into HTT_1-588_- YFP using EcoRV and NotI sites following the IDT protocol. Incorporation of point mutations were confirmed by DNA sequencing (MWG Eurofins).

### Caspase cleavage assays

As previously described, caspases were activated in HeLa cells expressing the indicated HTT_1-588_-YFP constructs by the addition of 1 μM staurosporine (STS) and 5 μg/mL cycloheximide (CHX) for the indicated times (9,16). Where specified, cells were transfected with the indicated constructs and treated with the indicated caspases inhibitors prior to the addition of STS/CHX. Typically, caspase cleavage of HTT_1-588_-YFP was detected in whole cell lysates. *In vitro* caspase assays were performed as previously described (9), using whole cell lysates. Briefly, lysates prepared from cells expressing FC or D552/586E 17Q-HTT_1-588_-YFP were aliquoted, resuspended in caspase cleavage buffer, and incubated with purified caspases for 1 h. Reactions were stopped by the addition of SDS Laemmli sample loading buffer.

### Pulse-chase analysis

Half-life of HTT_1-588_-YFP constructs were determined by pulse-chase analysis using Click chemistry, as previously described (17). HeLa cells were transiently transfected with the HTT_1-588_-YFP constructs using X-tremeGENETM^™^ 9 Transfection r agent (Roche), shortly after being seeded. 18 h post-transfection, cells were deprived of methionine and cysteine and incubated with 50 μM azidohomoalanine (AHA; Invitrogen) and non-labeled cysteine for 1 h (Pulse). After the 1 h pulse, the cells were washed with PBS and incubated with regular media (Chase). Cells were harvested after 0, 1, 2, 4, 8 and 24 h of chase, lysed in RIPA buffer and proteins were immunoprecipitated with goat anti-GFP polyclonal antibody (Eusera) and subjected to Click chemistry with Biotin-PEG4-alkyne (Sigma). Immunoblots for HTT1-588-YFP were conducted with rabbit monoclonal anti-GFP antibody (Eusera) and HTT_1-588_-YFP bio-orthogonally labeled with Biotin was detected with Alexa680 Streptavidin (Invitrogen).

### Filter-trap

Protein aggregation of insoluble HTT_1-588_-YFP was measured using a filter-trap assay, as previously described (18). After 48 h of transfection, HeLa cells expressing the indicated HTT_1-588_-YFP constructs were lysed in buffer containing 1% sarkosyl. Lysates were passed over cellulose acetate membrane using a vacuum manifold and probed by standard immunoblotting analysis with rabbit anti-GFP (Eusera) and detected by Licor.

### Quantification and Statistical analysis

Where indicated, Western blots were scanned using the Licor system. Bands were quantified using Image Studio V4.0. All statistical analysis was performed using Prism Graphpad v5.07. Half-lives were calculated by using a non-linear regression dose-response curve to determine the IC50. Where appropriate, one-way or two-way ANOVA was applied. Error bars are shown as standard error of the mean. Post-hoc tests are indicated where applicable. All experiments were performed 3 or more times.

## Results

### Blocking caspase cleavage at D586 promotes alternative cleavage at D552 and an unknown site

We previously noted that HTT_1-588_-YFP containing no mutations to block caspase cleavage, referred to as fully cleavable (FC), is cleaved predominantly at D586 by caspase-6 (9, 11). In order to confirm caspase cleavage of HTT at D586, we also previously generated a form of HTT_1-588_-YFP that is not cleavable at D586 (D586E) (9). These results are confirmed in Figure 1. The unknown band previously detected is indicated by an asterisk.

**Figure 1.**
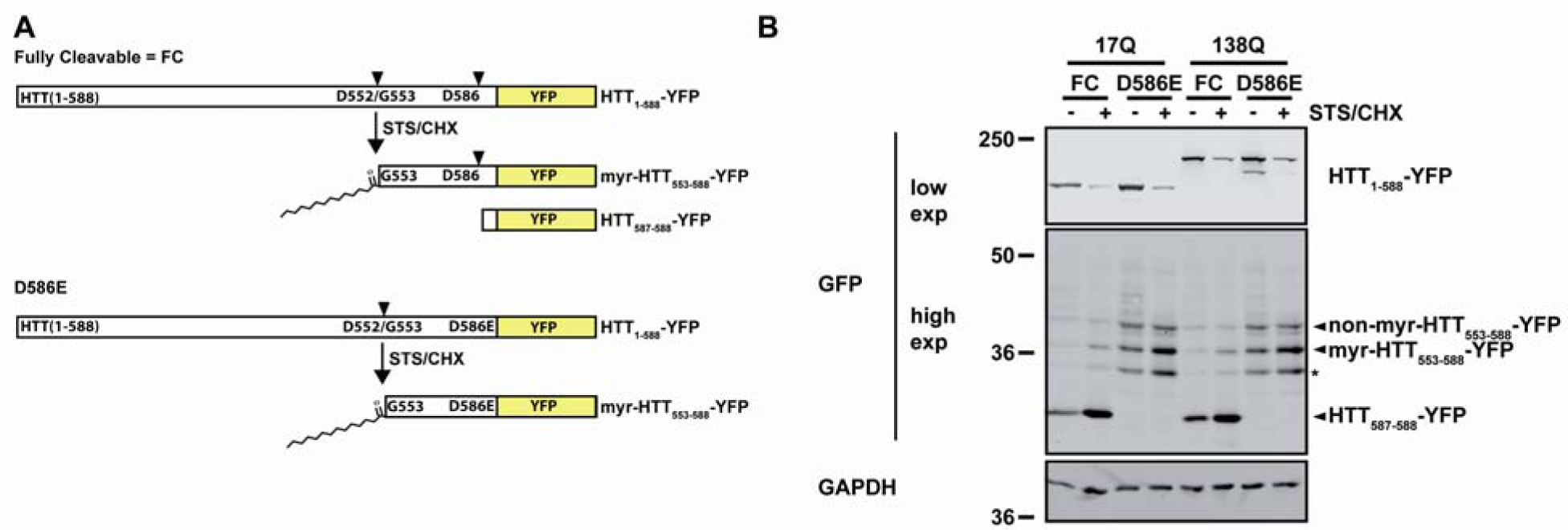
Blocking cleavage at D586 alters the proteolysis of HTT. **A.** Schematic representation of the predicted proteolytic fragments of caspase cleaved fully cleavable (FC, left) and D586E (right) HTT_1-588_-YFP. **B.** Caspase cleavage in HeLa cells expressing the indicated forms of wt (17Q) and mutant (138Q) HTT_1-588_-YFP was induced by adding 1 μM STS and 5 μg/mL CHX for 4 h after 20 h of transfection. Lysates were separated on 10% large SDS-PAGE gels and caspase-cleaved fragments were detected using anti-GFP antibodies. It was previously shown that the myristoylated and non-myristoylated HTT can be detected by mobility on SDS-PAGE. The D586 cleavage fragment has been previously described. Asterisk indicates the previously detected unknown band. Images are composites from the same gel.

As previously shown using click chemistry (11), identification of myristoylated and non-myristoylated HTT-YFP could be detected by SDS-PAGE mobility (Figure 2). Blocking caspase cleavage at D552 prevented the generation of 2 bands (Figure 2A) suggesting that both bands are related to the myristoylation status at G553 (11). Substitution of the essential glycine, required for myristoylation, to alanine or serine prevented the generation of only the lower of the two bands (Figure 2B), suggesting that it represents the myristoylated HTT fragment (myr-HTT_553-588_-YFP). Again, one band remained that migrated between D552 and D586 in the D552/D586E mutant. This band was also detected in fully-cleavable HTT_1-588_-YFP (Figure 1-3, 5 and 6) suggesting that it occurs naturally and is not due to alternative cleavage caused by inhibiting proteolysis at D586. The mobility of the unknown band between the bands generated by D552 and D586 caspase cleavage suggests that the unknown site is between the amino acids 552-586.

**Figure 2.**
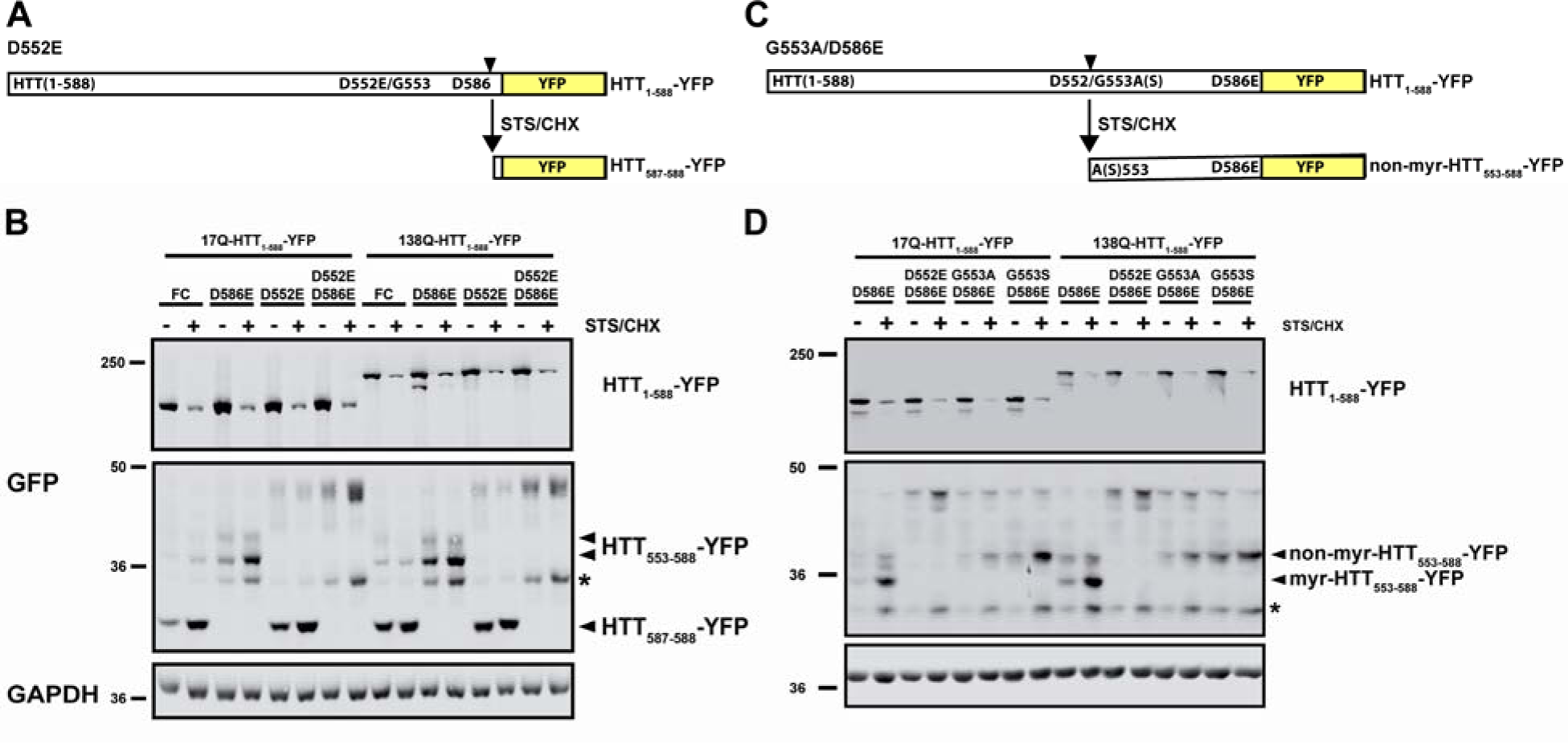
Cleavage of HTT at D552 generates 2 bands that correlate to myristoylation status. **A.**Schematic representation of the predicted proteolytic fragments generated by blocking caspase cleavage at D552 (D552E) in HTT_1-588_-YFP. **B., C.** Caspase cleavage in HeLa cells expressing the indicated forms of wt (17Q) and mutant (138Q) HTT_1-588_-YFP was induced by adding 1 μM STS and 5 μg/mL CHX for 4 h after 20 h of transfection. Lysates were separated on 10% large SDS-PAGE gels and caspase-cleaved fragments were detected using anti-GFP antibodies. * - denotes unknown cleavage product. Images are composites from the same gel.

**Figure 3.**
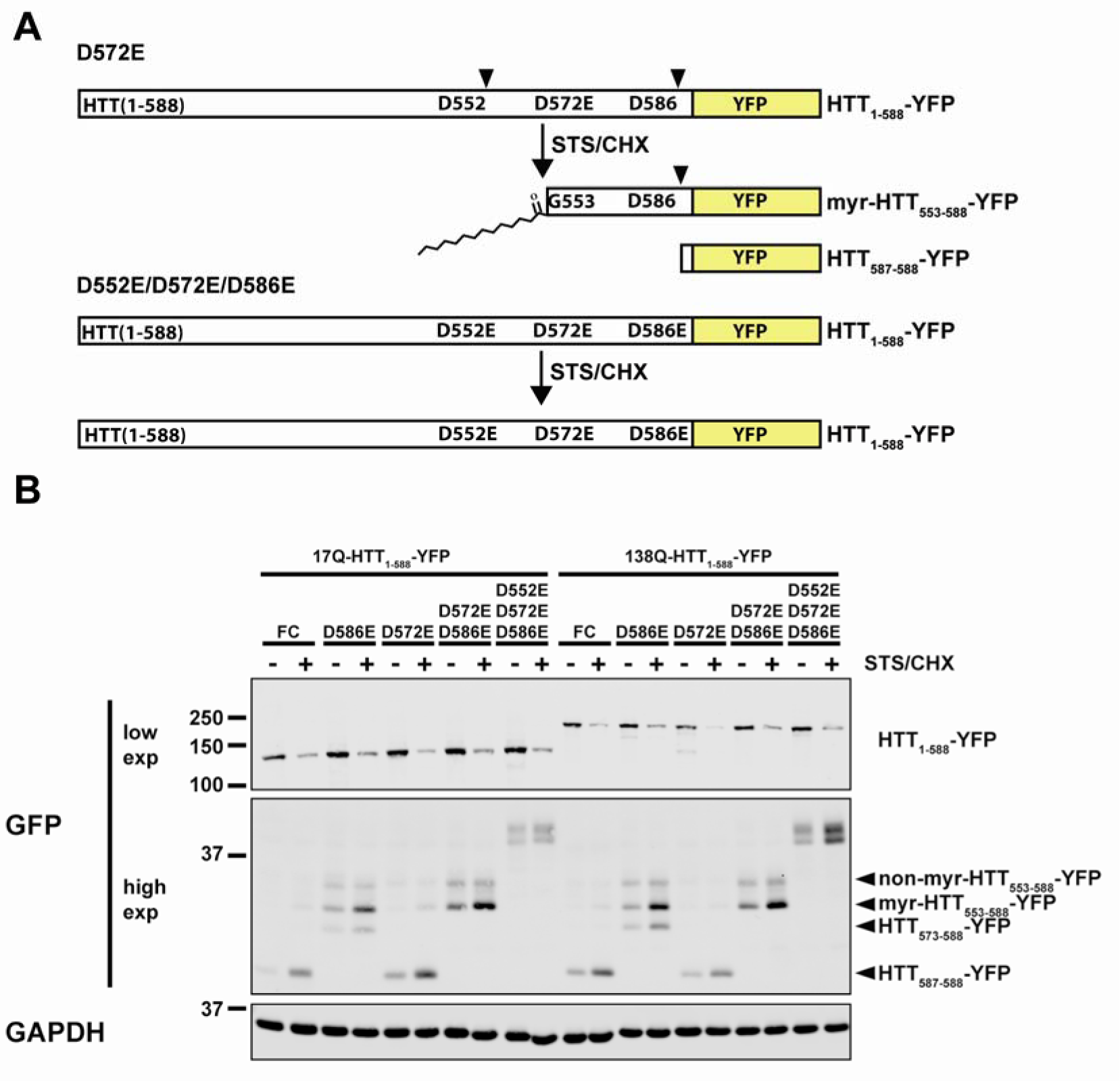
HTT is proteolysed at D572. **A.** Schematic representations of the predicted proteolytic fragments generated in (*Top*) FC HTT or (*Bottom*) by blocking caspase cleavage at D552, D572 and D586 in HTT_1-588_-YFP. **B.** HeLa cells expressing the indicated forms of 17Q and 138Q-HTT_1-588_-YFP bearing combinations of point mutations including D572E, D552E and D586E were treated with 1 μM STS and 5 μg/mL CHX for 4 h after 20 h of transfection to induce caspase cleavage. Lysates were separated on 10% large SDS-PAGE gels and caspase-cleaved fragments were detected using anti-GFP antibodies. The D572E mutation blocked the generation of the band that migrated between the caspase cleavage sites at D552 and D586. Images are composites from the same gel.

### HTT is proteolysed at D572

Subsequently, we sought to identify the unknown band. Based on the increase of the band intensity in the presence of the caspase inducers STS and CHX, we predicted that the generation of the band was due to caspases. Caspases proteolyse after aspartate residues (19). There are only two aspartate residues within the amino acid sequence between caspase cleavage sites D552 and D586, but only one site was predicted to be a caspase cleavage site using the caspase prediction analysis program CASVM (20), EGPD572. As confirmation, mutation of aspartate at D572 to glutamate (D572E) eliminated the detection of the new band in single (D572E), double (D572/D586E) and triple (D5552/572/586E) mutants (Figure 3B), thereby identifying that HTT_1-588_-YFP is proteolysed at D572.

### HTT is cleaved at D572 by caspase-1

In order to confirm that HTT is proteolysed by caspases during cell death with STS/CHX, cells expressing the double mutant D552/586E-HTT_1-588_-YFP, to specifically promote cleavage at D572, were treated with the irreversible pan-caspase inhibitor QVD-OPh. QVD-OPh completely inhibited cleavage at D572 in FC and D552/586E-HTT_1-588_-YFP (Figure 4B) confirming that the EQPD572 motif is a bona fide caspase-cleavage site.

**Figure 4.**
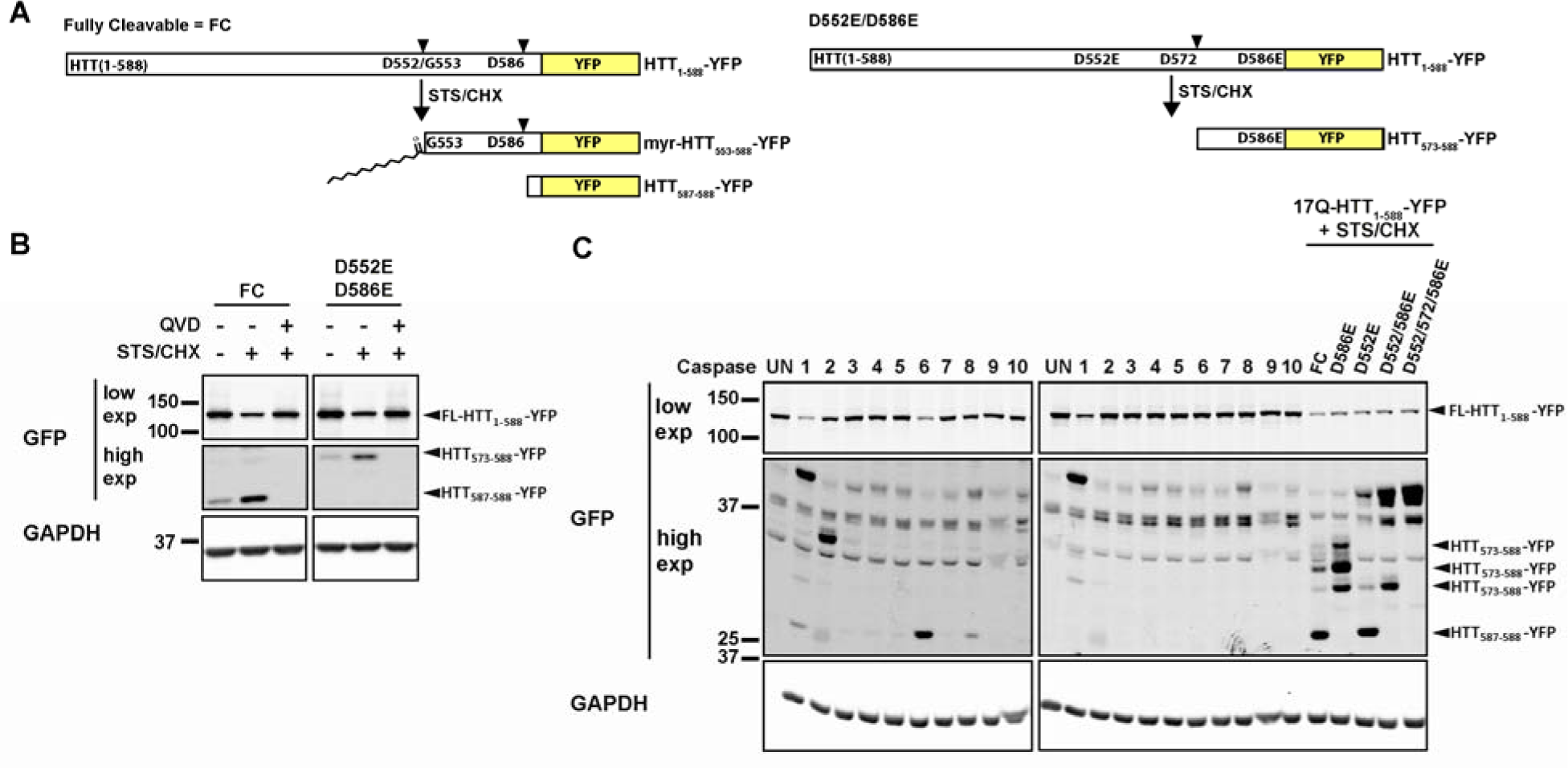
Cleavage of HTT at D572 is mediated by caspase-1. **A.**Schematic representation of the predicted fragmentation of FC and D586E-HTT_1-588_-YFP. **B.** HeLa cells transiently expressing 17Q FC-or D552/586E-HTT_1-588_-YFP were induced to undergo apoptosis, as described above, in the presence or absence of the general caspase inhibitor Q-VD-OPh. The band corresponding to proteolysis at D572 was not detected in the presence of the caspase inhibitor. **C.** Lysates from cells expressing 17Q FC-and D552/586E-HTT_1-588_-YFP were incubated with the indicated purified caspases. HTT_1-588_-YFP was cleaved at D572 by caspases 1 and 2.

Next, we sought to identify the caspase responsible for proteolysis at D572. FC-HTT_1-588_-YFP was used to detect caspase cleavage at D572 in unmodified HTT_1-588_-YFP, and D552/586E-HTT_1-588_-YFP was used to increase specificity for cleavage the D572. Both forms were predominantly processed by caspase-1 at D572, followed by, to a much lesser extent, caspase-2 (Figure 5).

**Figure 5.**
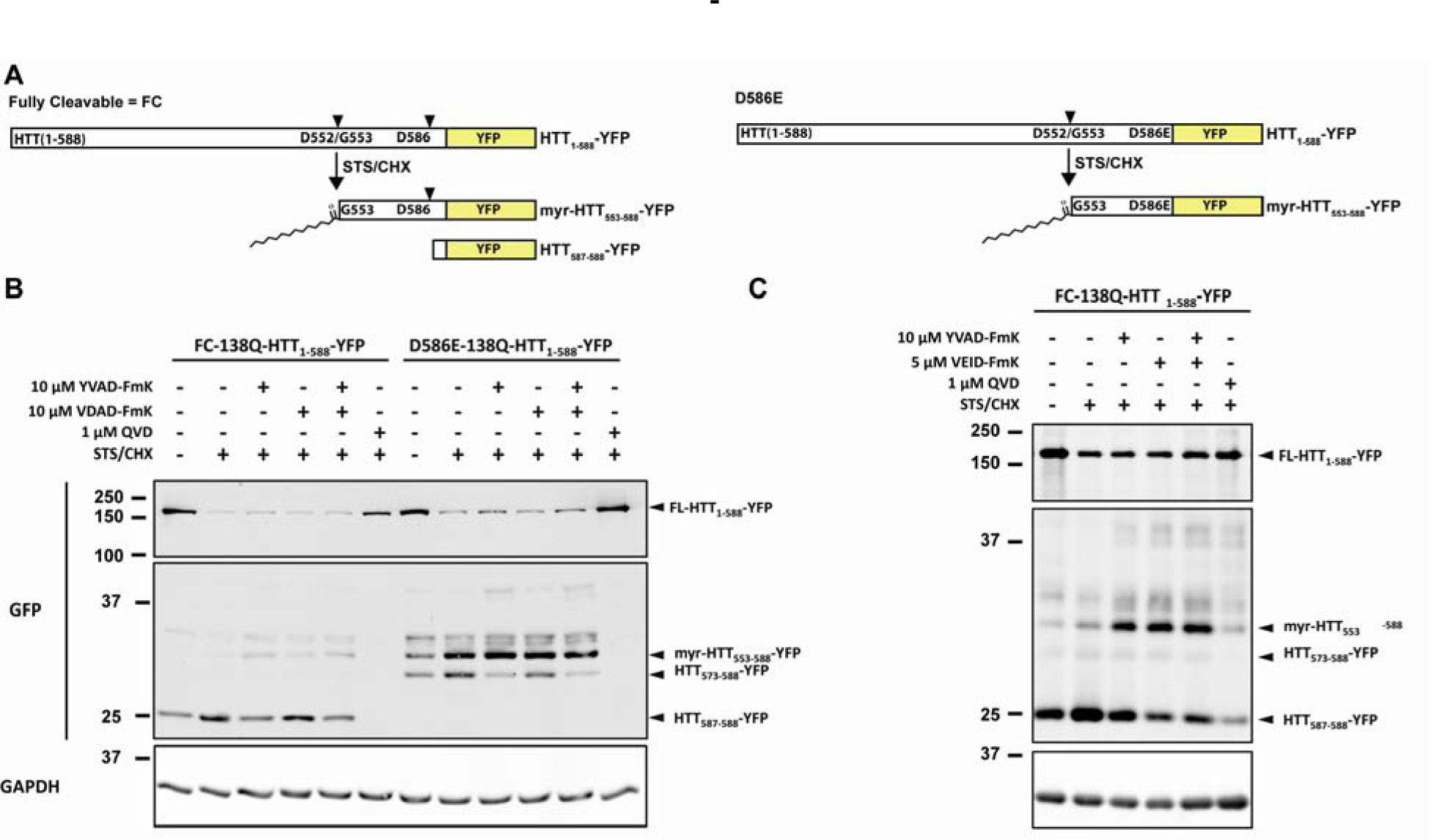
Inhibition of caspase-1 decreases cleavage at D572 and D586. **A.** Schematics representing the caspase cleavage profiles of FC and D586E HTT_1-588_-YFP. **B.** As described above, caspase cleavage at D572 in FC and D586E HTT_1-588_-YFP was followed in the presence or absence of 10 μM caspase-1 and 2 inhibitors (YVAD-FmK and VDAD-FmK, respectively) in the presence of STS and CHX. **C.** Caspase cleavage was induced with STS and CHX in the presence or absence of caspase-1 or 6 inhibitors (YVAD-FmK and VEID-FmK, respectively). Both inhibitors increased post-translational myristoylation at G553 through increased cleavage at D552.

### Inhibition of caspase-1 decreases proteolytic cleavage of HTT at D572 and D586 while promoting post-translational myristoylation at G553

In order to confirm proteolysis by caspases 1 and 2 at D572 in HTT, HeLa cells expressing FC-and D586E-HTT_1-588_-YFP, to promote cleavage at D572, were treated with specific caspase-1 and -2 inhibitors in the presence or absence of the cell stressor STS and CHX. As shown in Figure 5B, addition of the caspase-1 inhibitor YVAD-FmK noticeably decreased levels of caspase cleavage at D572, while addition of the caspase-2 inhibitor YDAD-FmK had more modest effects. Addition of the two caspase inhibitors appeared to decrease D572 cleavage to a greater extent.

Of particular note, the addition of caspase inhibitors, particularly caspase-1, appeared to increase post-translational myristoylation at G553 as well as decrease proteolysis at D586 in FC-HTT_1-588_-YFP (Figure 5B). Caspase-6 is the predominant caspase that cleaves HTT at D586 (9). Caspase-6 activity has also been shown to be activated by caspase-1 (21). Consequently, caspase cleavage of FC-HTT_1-588_-YFP was assessed in the presence of inhibitors of caspase-1 (YVAD-FmK) and caspase-6 (VEID-FmK) to specifically inhibit proteolysis at D572 and D586, respectively (Figure 5C). Caspase-1 inhibition decreased cleavage at D572 and D586. Caspase-6 inhibition only inhibited cleavage at D586. Notably, addition of either caspase inhibitor, or in combination, increased post-translational myristoylation at G553 and decreased proteolysis at D586.

### Blocking caspase-cleavage of HTT_1-588_-YFP at D572 decreases mHTT aggregation and increases soluble mHTT clearance

In order to determine how caspase cleavage at D572 regulates clearance of soluble mHTT, we sought to measure the half-life of HTT-YFP in the presence or absence of cleavage at the D572 site. Half-lives were determined using pulse-chase analysis and the methionine analog azidohomoalanine (AHA) (17). Strikingly, blocking caspase cleavage at D572 significantly reduced the half-life of both 17Q and and 138Q-HTT (2.3 h and 3.4 h, respectively) compared to their FC counterparts (3.8 h and 5.3 h, respectively).

The polyQ mutation of HTT is associated with increased aggregation of HTT (18). Consequently, we measured insoluble HTT to establish if the decreased half-life of soluble mHTT was not simply due to increased insoluble HTT aggregation. All forms of the 17Q-HTT_1-588_-YFP did not form appreciable aggregates. However, mutant 138Q-HTT_1-588_-YFP had significantly higher insoluble aggregates as expected. The decreased half-life associated with blocking caspase-cleavage at D572 only (D572E) was also associated with decreased insoluble mHTT aggregates (Figure 6C) suggesting that the decreased half-life is not due to increased aggregation of mHTT.

**Figure 6.**
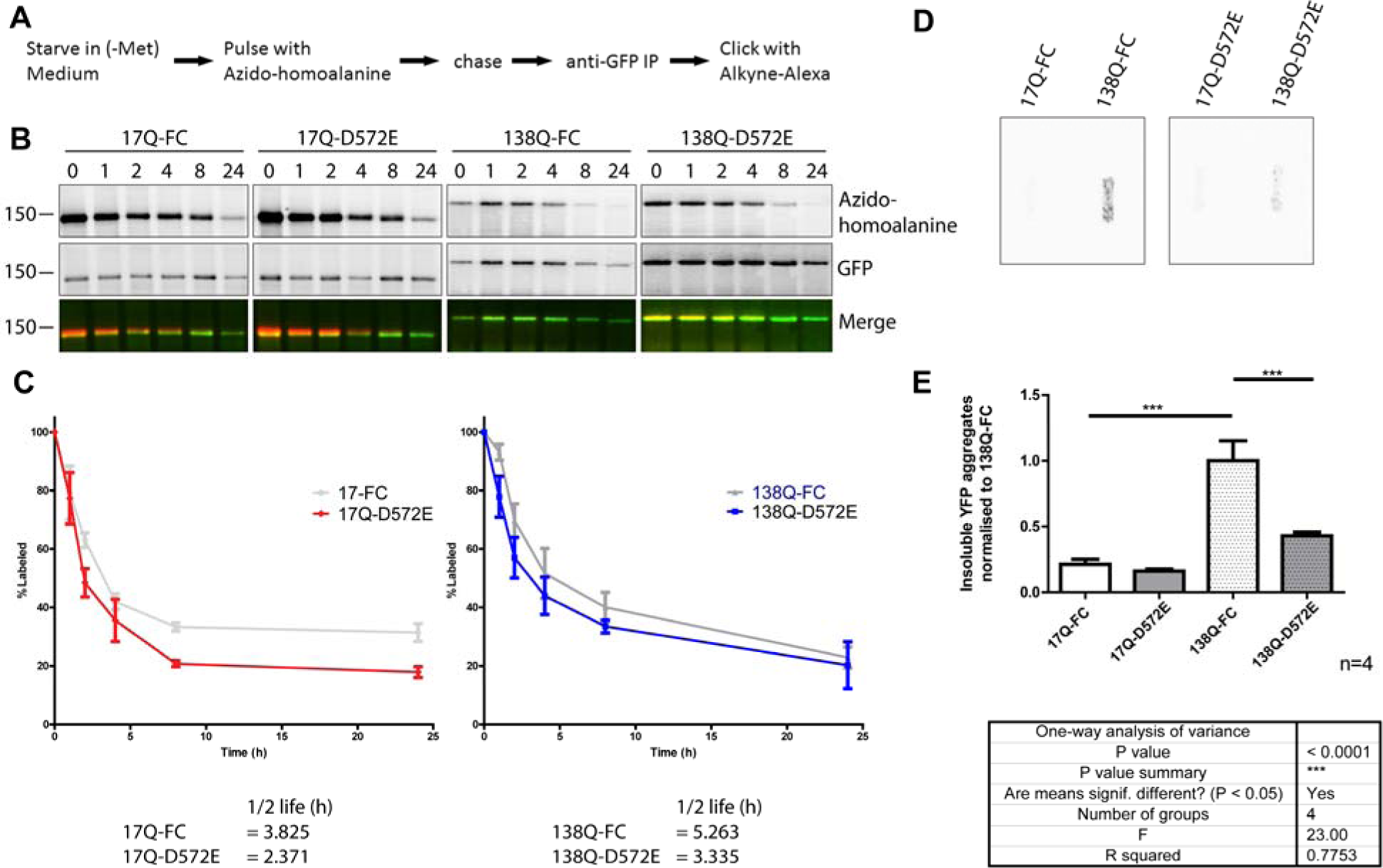
Blocking caspase cleavage at D572 increases soluble mHTT clearance and decreases insoluble mHTT aggregates. **A.** Schematic outlining the protocol for radioactive free pulse chase analysis using the methionine analog AHA. **B.** Western blot analysis of the indicated HTT_1_-588-YFP constructs immunoprecipitated from HeLa cells labeled with AHA and subjected to click chemistry for detection with streptavidin. C. Quantification of B presented as percent AHA label of GFP labeled compared to 0 h (or 100% labelled). **D.** Insoluble HTT_1-588_-YFP aggregates were detected using a filter-trap assay. Lysates were passed over a cellulose acetate membrane by vacuum and detected by immunoblotting. **E.** Quantification of insoluble HTT_1-588_-YFP aggregates detected in **E** are displayed as normalised to FC-13 8Q-HTT_1-588_-Y F P. Error bars represent SEM. *** - denotes significant values <0.0001.

## Discussion

Proteolysis of mHTT has been shown to be crucial in the pathogenesis of HD (8, 22). However, processing of HTT is a complex process mediated by numerous proteases, predominantly caspases and calpains (23). Previously, we have shown that proteolysis of mHTT at D586 is essential for the pathogenesis of HD (8, 9). In turn, we have recently identified alterations in the modification of HTT that can promote differential processing of HTT (11). For instance, the naturally occurring SNP leading to a missense mutation blocks both post-translational myristoylation at G553 and caspase-cleavage at D552. In turn, the G553E mutation also promotes proteolysis at D513 suggesting that there is co-operation or an inter-relationship with the PTM network of HTT in which some PTMs promote or inhibit others.

Herein, while studying the changes induced by blocking D586 cleavage, we identified a novel caspase-cleavage site. Furthermore, we identified the primary caspase responsible for cleavage at D572, namely caspase-1. In addition, inhibition of caspase-1 also inhibited proteolysis at D586; both indirectly and directly. Indirect inhibition was likely through inhibition of caspase-6 activation by caspase-1, as previously shown (21). Although caspase cleavage of HTT by caspase-1 has been shown previously (7, 9), specific cleavage of HTT at D586 by caspase-1 has not been shown.

Of note, inhibiting caspase cleavage of HTT at D572 (D572E) led to increased clearance of soluble mHTT (Figure 6), as well as a significant decrease in insoluble mHTT aggregates. This suggests that blocking cleavage at D572 is protective and that caspase-1 inhibitors may be viable targets for exploration in HD. Because caspase-1 can directly activate caspase-6 (21), inhibiting caspase-1 has the added benefit of inhibiting HTT cleavage by caspase-6 at D586, which is a critical event in the pathogenesis of HD. We predict that caspase-1 inhibition in HD may mediate a protective effect by preventing caspase cleavage of mHTT at D572 and D586 (Figure 7), thereby increasing the clearance of mHTT and preventing the formation of mHTT aggregates.

**Figure 7.**
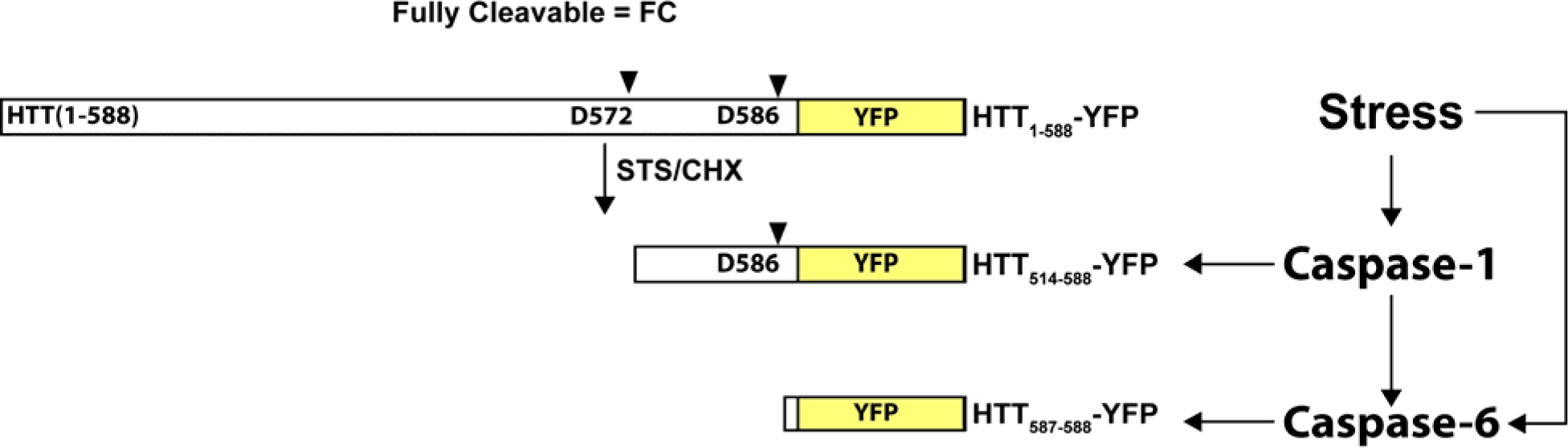
HTT is cleaved at D572 by caspase-1 upstream of caspase-6 mediated cleavage of HTT at D586. HTT is cleaved at D572 and D586 by caspases-1 and 6, respectively. Caspase-1 is upstream of caspase-6. Therefore, inhibition of caspase-1 decreases cleavage of HTT at both D572 and D586.

## ACKNOWLEDGEMENTS

The authors would like to thank Dr. Shaun Sanders for critical review of the manuscript and members of the Hayden laboratory for useful discussions. In particular, Dr. Niels Skotte was helpful in determining the half-lives of proteins. This work was supported by the Canadian Institutes of Health Research (CIHR 20R90174). DDOM was supported by postdoctoral fellowships from CIHR and the Michael Smith Foundation for Health Research as well as the Bluma Tischler Fellowship from UBC. MES was awarded a Vanier Canada Graduate Scholarship from CIHR.

